# Predictive E-prop: A biologically inspired approach to train predictive coding-based recurrent spiking neural networks

**DOI:** 10.64898/2026.02.12.705507

**Authors:** Davide Noè, Hideaki Yamamoto, Yuichi Katori, Shigeo Sato

## Abstract

The predictive coding framework offers a compelling model for temporal signal processing in the cortex. Recent studies explored its implementation in spiking architectures using Hebbian plasticity rules or offline learning; however, a biologically inspired model that enables gradient-based minimization of prediction errors remains an open challenge. In this work, we demonstrate that the predictive coding objective can be optimized using the online and local nature of the e-prop learning algorithm in recurrent spiking neural networks, creating the Predictive E-prop model. We demonstrate that the model is capable of learning complex time-series signals purely from self-supervised learning, using only its own prediction error as input, maintaining self-sustaining activity and reproducing the target’s underlying dynamics even in the absence of external stimuli. Furthermore, Predictive E-prop shows robust signal reconstruction abilities, effectively filtering noise and successfully interpolating sparse data. A comparative study against a backpropagation-based approach reveals that the two achieve comparable performance after training, confirming the viability of our model for timeseries generation tasks. These findings are particularly relevant for future developments in neuromorphic hardware, offering a purely self-supervised, gradient-based model that could provide significant advantages in power efficiency and computational ability.

## 1. Introduction

The mammalian brain is not a simple input-output device but instead possesses internal models that generate predictions of the environment and shape sensory processing [1]. Since its inception as a theory of visual perception [2], such mechanism of cortical information processing, i.e. predictive coding, has been generalized as a computational framework and demonstrated to be implementable in recurrent neural network (RNNs) models [3, 4]. Unlike traditional RNNs driven primarily by forward signal propagation, predictive coding-based RNNs utilize error-driven signaling to intrinsically suppress redundant activity, thereby improving computational efficiency and reducing energy consumption [5, 6]. The objective of learning thus becomes minimizing the prediction error by optimizing the internal model of the environment generated purely from “perception”, eliminating the requirement for external, human-annotated data [7]. The current landscape for predictive coding-based RNNs is very rich, with implementations ranging from reservoir computing approaches based on rate-coding [8] and spiking [9] neural networks, to more biologically plausible Hebbian learning-based ones [10, 11]. However, an online and local gradient-based approach to minimize prediction error remains an open challenge.

Methods based on gradient descent are particularly interesting, because they have shown exceptional representation learning abilities, allowing models to learn complex, non-linear representations directly from raw data [12, 13]. The most prominent approach for training RNNs is Backpropagation Through Time (BPTT), a gradient descent based algorithm that operates by unraveling the recurrent network over the time dimension and then performing backpropagation, treating each time-step as a layer in a deep feedforward network [14]. Although highly effective, BPTT suffers from memory costs that grow linearly with sequence length, making it impractical for nontrivial time-sequences [15]. Furthermore, its weight update occurs after all data has been processed and requires complete information from the entire network, which is in stark contrast to the local, online nature of biological neural networks [16].

A biologically plausible, gradient-based algorithm, eligibility propagation (e-prop), has been proposed to tackle these challenges [17]. The e-prop learning algorithm employs surrogate gradient methods [18] to allow for the computation of gradients in recurrent spiking neural networks (RSNNs). The core mechanism of e-prop is a form of neo-Hebbian three factor learning [19]; neurons accumulate eligibility traces of their activity while processing information; these traces are then activated by a “third factor,” a neuron-specific learning signal that triggers synaptic plasticity according to the amount of traces present. This local approach has proven to lead to competitive performance in noisy environments and sparse networks [20], removing the resource overhead of BPTT. The power efficiency of e-prop has also been validated by its implementation in neuromorphic hardware, including general-purpose chips, such as Loihi [21] and SpiNNaker 2 [22] as well as in ASICs such as ReckOn [23]. However, its integration in predictive coding architectures remains unexplored, largely because prior research focused on optimizing eprop for temporal credit assignment and predictive coding for hierarchical inference, leaving the potential synergy between their local learning mechanisms under-examined.

In this work, we bridge this gap by integrating the e-prop algorithm with predictive coding-based RSNNs. We term the resulting model ‘Predictive Eprop’, emphasizing its role as a learning principle rather than a task specific model. In Predictive E-prop, the neuromodulatory “third factor” described above is derived strictly from the local prediction error rather than a task-specific external signal. To evaluate this approach, we test the performance of our network on two distinct dynamical systems—a periodic limit cycle (sine wave) and a 3D chaotic strange attractor system (Lorenz attractor)—across three distinct tasks. Our results show that the predictive coding objective can be optimized using the e-prop algorithm, allowing the network to adapt to target dynamics and maintain self-sustaining activity without external input. The network also showed robust reconstruction abilities, recreating the correct signal despite significant amounts of sub-sampling and filtering out high amounts of white noise. Lastly, comparing Predictive E-prop’s generative ability with an predictive-coding RSNN trained with truncated BPTT (TBPTT), we observed that the two algorithms converged to comparable performance, underscoring the suitability of Predictive E-prop for signal forecasting applications.

## 2. Methods

### 2.1 Network model

The structure of the model used in this work comprises four functional weight matrices; an input (*W* ^in^) and feedback (*W* ^fb^) matrices, which are kept static and defined as 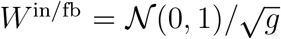, where *g* is a tunable input gain parameter, and a recurrent (*W* ^rec^) and output (*W* ^out^) matrices, which undergo training and are initialized as 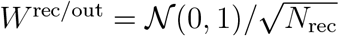, where *N*_rec_ is the number of neurons in the recurrent layer (Fig. 1A).

**Figure 1.**
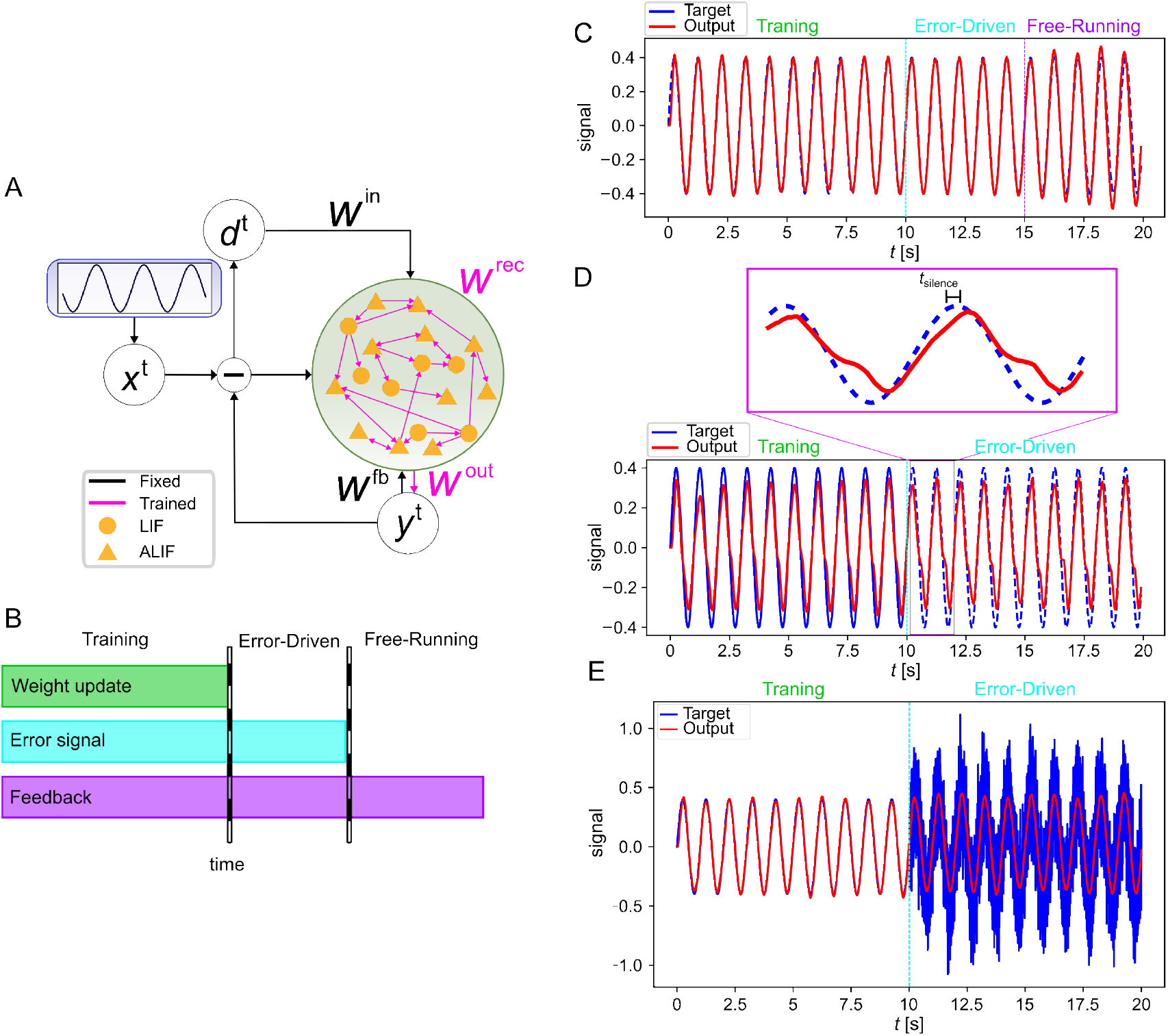
A) Structure of the network used in the experiment; the fixed connections are represented in black while the evolving weights are highlighted in magenta. B) The three phases of the experiment: training, error-driven, and free-running. C) Time series generation task on a sine wave input signal. D) Representation of the time-series reconstruction task; during the error-driven phase the input signal is provided every *t*_silence_ time-steps. E) Noise filtering task on a sinusoidal target (dashed).

Following the architectural principles of Long-Short-Term Memory SNNs (LSNNs) [24], our recurrent layer comprises 300 neurons divided into a combination of *N*_LIF_ leaky-integrate-and-fire (LIF) and *N*_ALIF_ adaptive LIF (ALIF) neurons [25]. More information about the make-up and initialization of the network for different experiments can be found in Tables SI to SIV in the Supplementary Materials. The internal state of a neuron *i* at time *t* is defined as 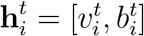, where 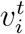 is its membrane potential and 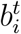 is its adaptive firing threshold. The membrane potential evolves by integrating synaptic inputs, feedback signals, and stochastic noise according to:

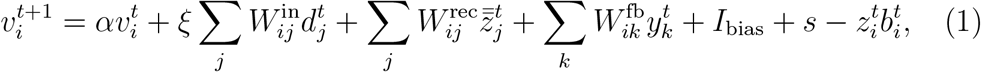

where 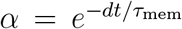 is an exponential decay term, with time step *dt* and membrane time constant *τ*_mem_.

Here, the output of the network 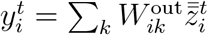, with 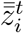 being the spike vector passed through a double exponential filter (see Eq. 2), represents a generative prediction and the input signal corresponds to the prediction error 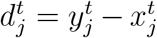, where 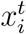 is the target signal. The network also receives its own prediction through the feedback layer to stabilize neural dynamics [26]. The *I*_bias_ term represents a bias current, and *s* is an Ornstein-Uhlenbeck noise term [27] with mean, standard deviation, and time-constant of *µ*_s_, *σ*_s_, and *τ*_s_, respectively. *ξ* = {0, 1} is a constant that controls the error input, allowing the network to transition from the error-driven to the free-running phase.

The firing threshold is defined as 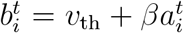, with 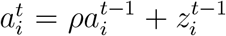. Here, 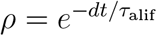 with *τ*_alif_ being the time decay constant for the adaptable threshold. *β* is set to a heuristically determined constant for ALIF neurons and to 0 for LIF neurons. The last term on the RHS of Eq. 1 implements a repolarization mechanism, reducing the membrane potential by the threshold value 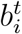 after a spike is emitted. A refractory mechanism is also implemented, keeping the membrane potential fixed for 5 ms after the repolarization. The spikes are computed using a Heaviside step-function *θ* as 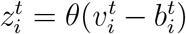 and then passed through a double exponential filter defined as:

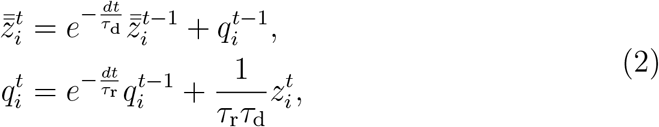

with 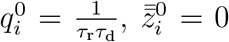, and *τ*_r*/*d_ time constants for rise/decay, respectively. The values of each parameters are listed in Tables SI–SIV of the Supplementary Material.

### 2.2 Learning algorithm

The weights in the recurrent layer *W* ^rec^ and the output layer *W* ^out^ are trained using the e-prop learning algorithm [17] and a standard gradient descent method, respectively. All other weights are kept fixed throughout the experiment.

For the recurrent layer, e-prop is implemented following:

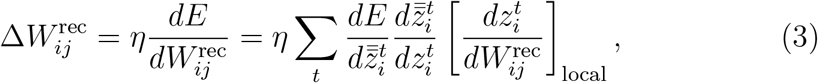

where *E* defines the objective of predictive coding, corresponding to the minimization of prediction errors and *η* is the learning rate:

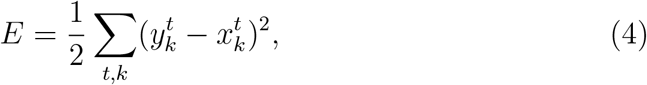

where 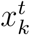 and 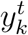 are the sensory input and prediction output, respectively, for the *k*-th dimension at time *t*. The first term on the RHS of Eq. (3) is the learning signal *L*, which corresponds to the local gradient of the objective of predictive coding with respect to the neural activity, and takes the form:

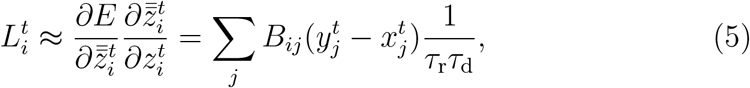

where *B*_*ij*_ is a fixed random matrix used to implement broadcast alignment [28]. The last term in Eq. (3), called the eligibility trace *e*_*ij*_, is computed as:

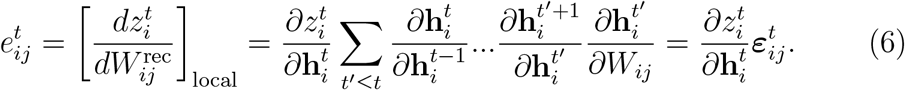

Since the spike function 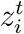 is not differentiable, the derivative 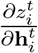 is approximated by a tent function:

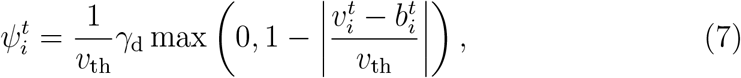

where *γ*_*d*_ is a constant that controls the height of the tent function. The eligibility vector ***ε***_*ij*_ = [*ε*_*ij,v*_, *ε*_*b,ij*_] in Eq. (6) can be computed recursively according to:

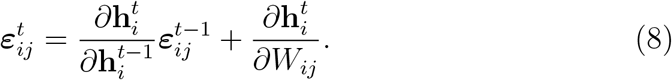

For LIF neurons, ***ε***_*ij*_ is just the presynaptic spike train passed through a low-pass filter, defined as 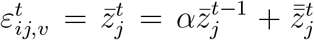. For ALIF neurons, the adaptable threshold requires a more careful treatment, giving the following contribution to the eligibility vector:

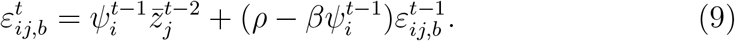

In summary, the final form of the eligibility trace is as follows:

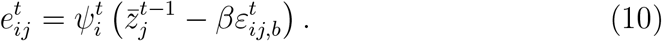

See refs. [17, 20] for a full derivation.

The evolution of the output layer weight from neurons *j* to *i*, 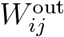, is computed as follows:

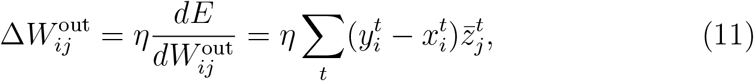

where 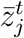 is the low-pass-filtered spike train defined above, 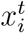 is the target signal, and 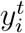 is the network’s prediction.

The network evolution occurs every *t*_delay_ ms to accommodate a homeostatic plasticity mechanism added to the loss function to homogenize neural activity. A weight decay term was also implemented to prevent excessive weight growth. For further information on both regularization terms, see Supplementary Materials “Network regularization.”

### 2.3 Experiment structure

An experiment consisted of repeated epochs, each comprising three separate phases (Fig. 1B):

- Training phase: The network receives input signals. The recurrent and output weights are evolved according to Eqs. (3) and (11), respectively.
- Error-driven (recognition) phase: All weights in the network are fixed. The input signal is still provided.
- Free-running (generation) phase: The network no longer receives the input signal and is required to generate sustained activity from its own feedback and recurrent layer activity.

Details about the length of each phase are provided in the Supplementary Materials, Tables SI to SIV.

The total number of epochs in an experiment was set to 50, unless otherwise specified. The weights were initialized at the start of the experiment as defined in Sec. 2.1 and carried over through successive epochs. Successful behavior during the free-running (generation) phase was used as evidence that the network has achieved an internal generative model.

### 2.4. Tasks

In this work, the network was tested on three different tasks: time-series generation, time-series reconstruction, and noise filtering. These three tasks have been chosen because they probe complementary aspects of predictive coding, namely its ability to create an internal generative model, the stability of the model under information gaps, and its resilience to overfitting, respectively.

The objective of the time series generation task (Fig. 1C) was to learn and recreate the internal dynamics of the original dynamical system, sustaining this activity even without receiving any feedback error during the free-running phase.

In the time-series reconstruction task (Fig. 1D), the model underwent only the training and error-driven phases. During the training phase, the network received the full prediction error; during error-driven phase, the feedback signal was sent only every *t*_silence_ ms. The model was tasked to learn and sustain the correct activity, outputting the correct signal despite the sub-sampling.

The noise filtering task (Fig. 1E) was designed similarly to the reconstruction task, with the network receiving the normal feedback error during the training phase and a noisy signal during the error-driven phase. Random, uncorrelated noise was added to the input signal using a Gaussian process:

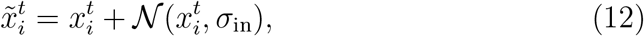

where 𝒩 represents a random normal distribution with mean 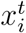 and SD *σ*_in_. The resulting feedback error during the error-driven phase was then given by:

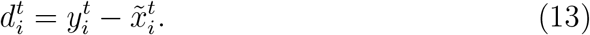

For all three tasks, two time series signals were considered, i.e., a sinusoidal signal corresponding to a single point attractor and a chaotic Lorentz attractor signal. The function generating the sine wave is:

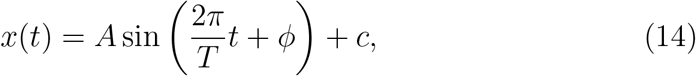

where *A* is the amplitude, *T* is the oscillation period, *ϕ* is the phase, and *c* is an offset. The parameters for the sine wave are summarized in Tables SI and SIII in the Supplementary Materials.

The time evolution of the Lorenz attractor is described in the following equations:

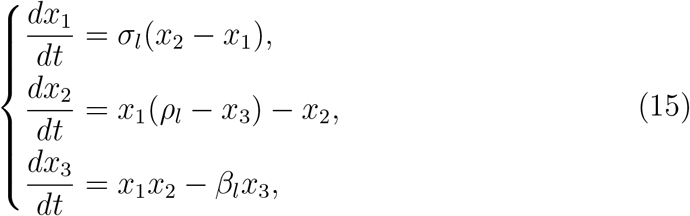

where *σ*_*l*_ = 10, *ρ*_*l*_ = 28 and *β*_*l*_ = 8*/*3. The Lorenz attractor data has been rescaled using a factor Γ_*l*_(= 100) for better tractability. The parameters used to generate the Lorenz attractor are provided in Tables SII and SIV in the Supplementary Materials.

Dynamic time warping (DTW) [29] was used to quantify network performance, as its ability to decouple phase and amplitude shifts from signal divergence makes it a preferable choice over the root mean square error measure for this type of task. Defining a warping path **Φ** = (Φ_*x*_, Φ_*y*_) with Φ_*x/y*_ mapping the elements of the prediction *y* and the target signal *x* to a common time axis of length *K*, the DTW distance is computed as the cumulative cost over all possible paths:

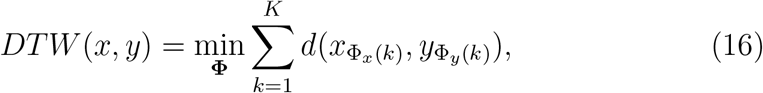

where *d*(·) is the Euclidean distance, and 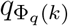 is quantity *q* = {*x, y}* warped by its relative warping function at time-step *k*.

## 3. Results

Predictive E-prop enables RSNNs to autonomously generate structured temporal patterns while retaining robust performance under severe signal degradation and noise. First, a signal generation task is used to assess the network’s ability to represent and maintain the attractor landscape of the target signal [30]. Building on this, we tested the network on a reconstruction task, probing the model’s ability to leverage its internal state to bridge information gaps [31], effectively simulating more realistic situations such as limited sampling rates in sensors or packet loss during communication. We then test the model on a noise filtering task to check that the network is not overfitting the input, but is learning the underlying dynamics of the target system [32]. Lastly, to contextualize the results obtained with our model, we compare its generative abilities against a backpropagation-based approach.

### 3.1. Time-series generation

As an initial benchmark, the generative ability of the network has been tested using a sine-wave time-series (see Fig. 1C). As shown in Fig. 2A, the network was successfully trained to self-sustain coherent spiking activity even in the absence of external stimuli, achieving a comparable DTW distance with respect to the other two phases during the free-running phase (Fig. 2B). Consistent with this result, the firing rate during the free-running phase evolved to match the training and error-driven phases (Fig. 2C).

**Figure 2.**
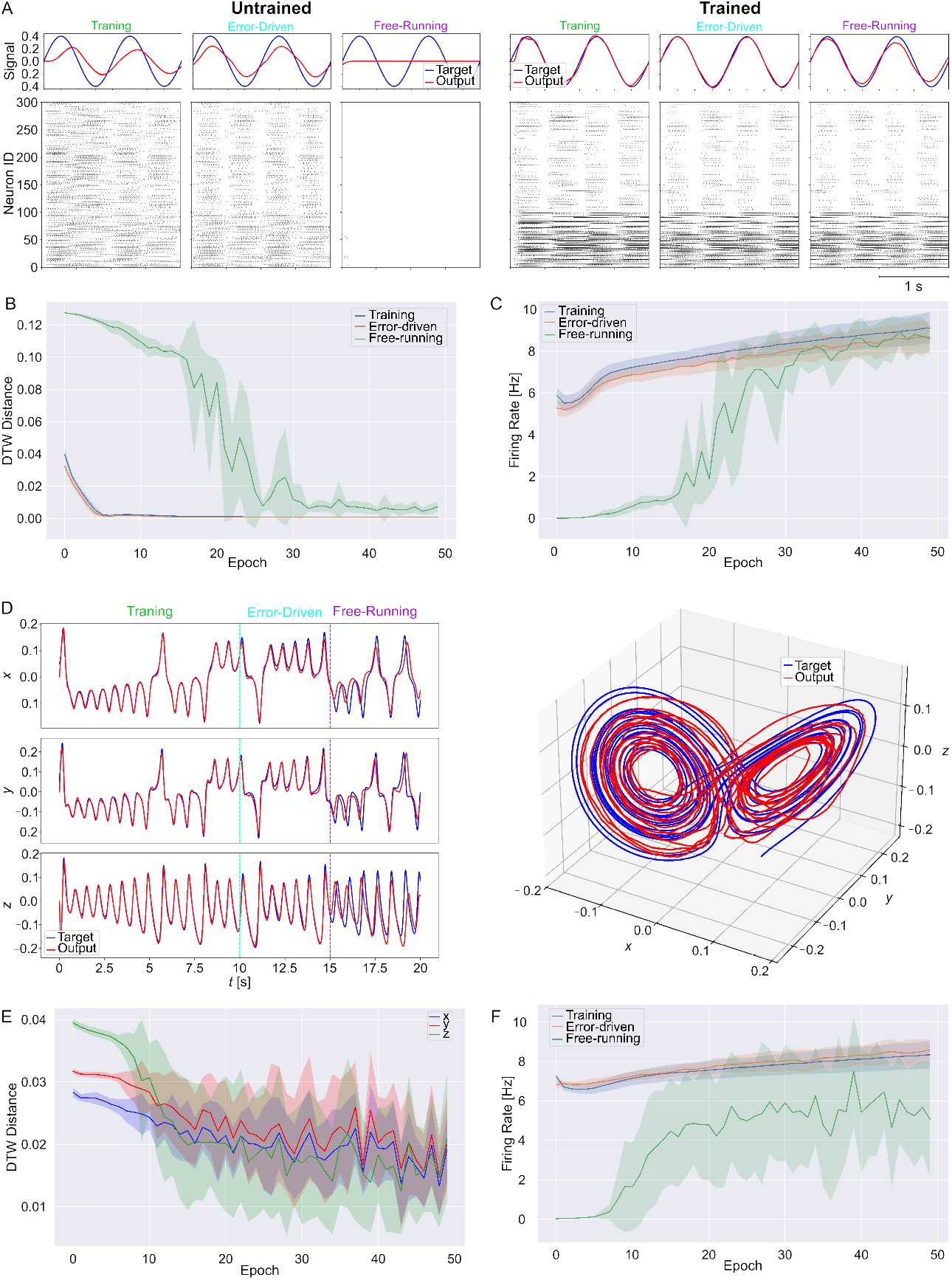
A) Examples of neural activity during the sine wave generation task across the three different phases for an untrained (left) and trained networks (right). B) Evolution of the DTW distance across the three phases of the experiment. C) Average firing rate during the training (blue), error-driven (orange) and free-running (green) phases. D) Example of signal generation for the Lorenz attractor projected on the three spatial dimension (left) and in 3D (right). E) DTW during the free-running phase for the three spatial dimensions throughout the epochs, highlighted in bold blue (*x*), red (*y*) and green (*z*). F) Average firing rate during simulation for the Lorenz attractor task. Panels B, C, F, and G represent the average and standard deviation over 10 randomly initialized runs.

The stable convergence was dependent on the homeostatic plasticity mechanism to homogenize activity alongside a temporally correlated Ornstein-Uhlenbeck noise to maintain neuron excitability. Without these factors, while the model successfully converged to the target dynamics during the training and error-driven phases, it experienced either runaway, synchronous firing or the complete collapse of neural activity after the transition to the free-running phase.

To further assess the network’s ability to learn and reproduce the target’s dynamics, we repeated the time series generation task on the Lorenz attractor dataset (Fig. 2D). Transitioning from a 1D periodic sine wave to a 3D chaotic attractor presented a significant challenge. Using the hyperparameter configuration set for the previous task led to a complete drop in neural activity when transitioning from the error-driven to the free-running phase. A fine-tuning of the static input and feedback layer gain parameter (*g*) was necessary for the network to achieve the required self-sustaining dynamical regime (see Table SII). With this new configuration, the network correctly learned to reproduce the Lorenz attractor dynamics (Fig. 2D), with a significant decrease in DTW distance across all three dimensions during the free-running phase (two-sided Welch *t*-test, *p <* 0.05, *N* = 10) (Fig. 2E). We also confirmed that the firing rate stabilized around the same value achieved during the training and error-driven phases without external stimuli for this target signal as well (Fig. 2F).

### 3.2 Time-series reconstruction

One line of evidence for predictive information processing in the cortex is the perception of subjective contour in, e.g., Kanizsa triangle [33, 34], in which observers perceive well-defined edges in regions of uniform luminance as a result of contextual inference. Related forms of predictive completion have also been observed in temporal domains, including auditory perception [35] and motor control [36]. To test whether an analogous predictive completion mechanism could emerge in the Predictive E-prop model, the network was tasked to reproduce a sine wave signal despite different levels of sub-sampling.

When the input signal was made sparse by muting the signal every *t*_silence_ ms, there was a significant increase in DTW distance at higher values of *t*_silence_ (Fig. 3A). Despite this performance deterioration, the network remained consistent in generating the sine wave signal from the intermittent error feedback (Fig. 3B). Achieving this stability required a fine-tuning of the homeostatic regularization intensity (Table SIII). By lowering the strength of the homeostatic regularization term, we prevented it from interfering with the error signal, maintaining the network’s susceptibility to the sparse error feedback. Consequently, the firing rate remained stable around the same value across all sub-sampling levels (Fig. 3C).

**Figure 3.**
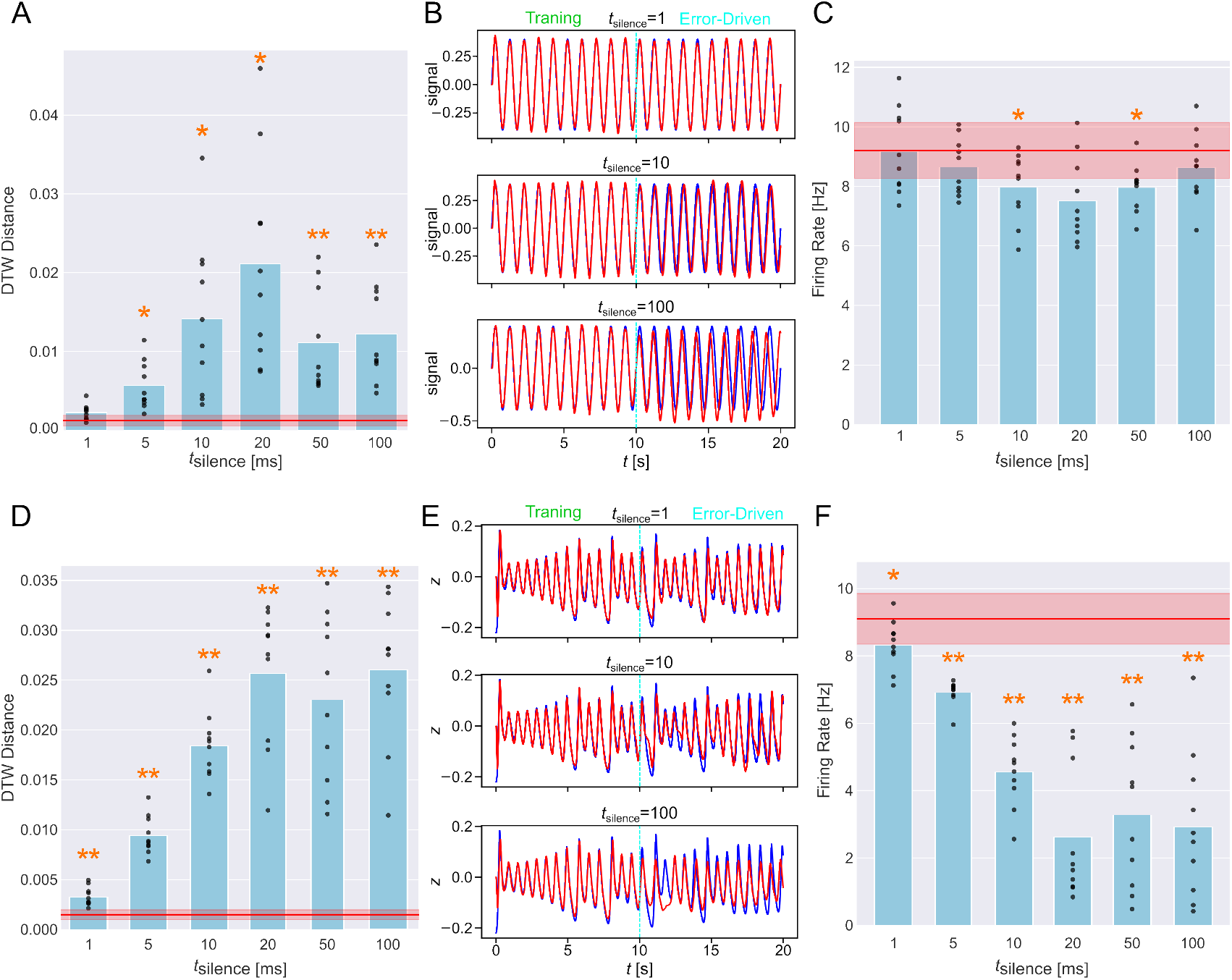
A, D) Average DTW distance during the error-driven phase for different levels of *t*_silence_ in the sine wave task (A) and the Lorenz attractor task (D). For the Lorenz attractor task, the values are for the *z*-dimension. B, E) Examples of signal reconstruction on the sine wave (B) and the Lorenz attractor (E), for *t*_silence_ = 1 ms, *t*_silence_ = 10 ms, *t*_silence_ = 100 ms. C, F) Average firing rate during the error-driven phase at different levels of sub-sampling for the sine wave task (C) and the Lorenz attractor task (F). In panels A, C, D, and F, points represent the average over the final ten epochs of each of the ten randomly initialized runs, while bars represent the average across the samples. In these panels, the red lines and shaded areas denote the mean and standard deviation, respectively, for data at *t*_silence_ = 0 ms. The asterisks denote p-values computed against the baseline: \**p <* 0.05 and \*\**p <* 0.01, two-sided Welch *t*-test (*N* = 10).

To further investigate the reconstruction ability of Predictive E-prop, we extended the time-series reconstruction task to the Lorenz attractor system. For this task, the interval between weight updates (*t*_delay_) was increased from 10 ms to 100 ms. This effectively increased the temporal window for firing rate regularization, allowing the network to ignore short-term fluctuations while preventing long-term quiescence, showing a fundamental balance between stability and preserving the complex target dynamics.

We found that the model’s performance was significantly more sensitive to the temporal sparsity of the feedback (Fig. 3D). Specifically, the network became incapable of accurately reconstructing the target chaotic trajectory above *t*_silence_ = 10 ms (Fig. 3E). Studying the network’s dynamics, we observed that this failure was tied to a collapse in neural activity, with the firing rate significantly dropping at higher values of *t*_silence_ (Fig. 3F). Nevertheless, the model was capable of reconstructing the three-dimensional, chaotic signal up to moderate levels of sub-sampling, indicating robustness of predictive reconstruction despite sparse feedback.

### 3.3 Noise filtering

Another characteristic feature of predictive coding is the ability to filter out noise from the input signal by referencing incoming signals against an internal model [8]. To examine this capacity in the current framework, we investigated whether Predictive E-prop could reconstruct a clean target from a noisy signal. The experimental design followed the signal reconstruction task described in Section 3.1 except that the error-driven (recognition) phase was tasked with a noisy signal and that there was no free-running (generation) phase. During the training phase, the model received the target signal without noise. The noise was modeled with a Gaussian process centered around the signal with tunable standard deviation *σ*_in_. The feedback error during the error-driven phase was then computed using the noisy signal.

Testing on the sine wave, we found that the network was capable of mitigating significant amounts of noise (Fig. 4A). Its performance remained stable up to *σ*_in_ = 0.2 (two-sided Welch’s *t*-test, *p >* 0.05, *N* = 10). Despite performance decreasing significantly at *σ*_in_ = 0.5, the error with respect to the target signal’s amplitude was found to be less than 3%, proving the robustness of the model to noise (Fig. 4B). The network’s firing rate at the end of the error-driven phase stabilized around 9 Hz consistently across noise levels, with the same pattern of degradation observed in the DTW distance (Fig. 4C).

**Figure 4.**
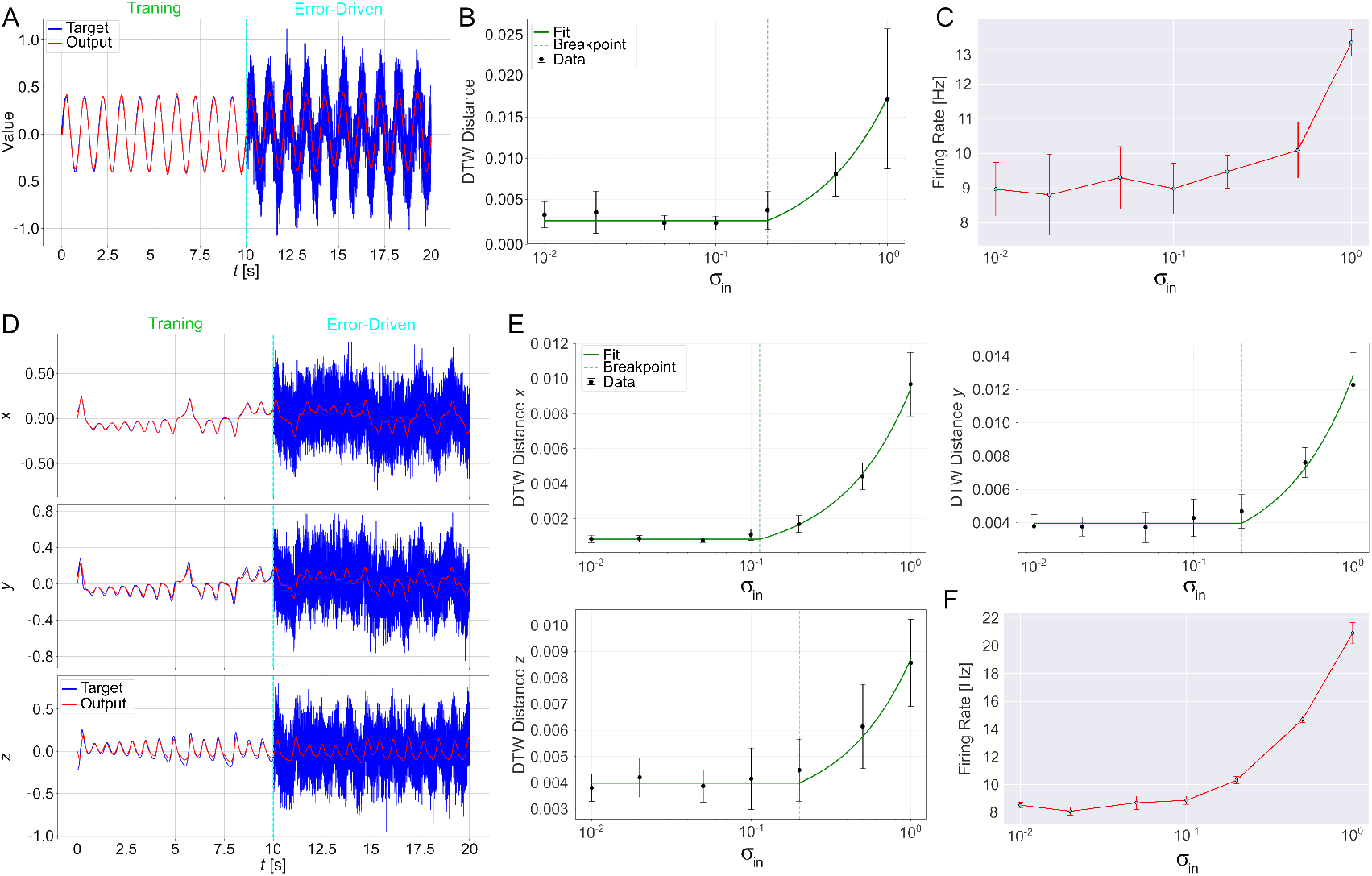
A, D) Example of noise filtering task for the sine wave with *σ*_in_ = 0.5 (A) and the Lorenz attractor with *σ*_in_ = 0.2 (D). The target and output signals are shown in blue and red, respectively. B, E) Average DTW distance during the error-driven phase in the final ten epochs at different noise levels for the sine wave (B) and the Lorenz attractor (E) tasks. C, F) Average firing rate over during the error-driven phase in the final ten epochs at different noise levels for the sine wave (C) and the Lorenz attractor (F) tasks. Panels B, C, E, and F represent the mean (dot) and standard deviation (error-bar) computed over ten randomly initialized runs. The data in Panels B and E were fit with piecewise regression using a constant line until the detected threshold and a linear equation afterwards.

A qualitatively similar trend was observed for the Lorenz attractor task, indicating that the noise-filtering behavior of the model extends beyond simple periodic dynamics (Fig. 4D). As summarized in Fig. 4E, the network consistently outputted the correct signal with noise up to *σ*_in_ = 0.2, above which a linear degradation in performance was observed. We again analyzed the average firing rate of the network in Fig. 4F, confirming the trend observed in the DTW distance. The firing rate remained consistent under *σ*_*in*_ = 0.2, supporting the hypothesis that the network is generating a robust, self correcting internal model despite the stochastic interference.

### 3.4 Comparison with truncated BPTT

Finally, to better contextualize our results in the landscape of RSNN training, we compared the performance of Predictive E-prop against an RSNN trained using TBPTT [37] under the same prediction-error objective, isolating the effects of the learning rule. We chose TBPTT because of its ability to mitigate the exploding gradient problem [38] and to handle a constant stream of data [39], allowing for a more fair comparison with our model. The model used for the two simulations have been kept the same, only the training algorithm has been changed. For more details on TBPTT see Supplementary Materials Section “Truncated BPTT”. The truncation interval for TBPTT was set to *k* = 100 ms, which was found to be optimal for the task at hand (see Fig. S1B in Supplementary Materials).

Experiments on the sine wave generation task revealed that the two models achieved comparable performance (Fig. 5A). As shown in Fig. 5B, the evolution of the DTW distance during the free-running phase converged to the same target performance after training (two-sided Welch’s *t*-test, *p >* 0.05, *N* = 10). These results corroborate the viability of Predictive E-prop as a functional alternative to backpropagation based approaches, demonstrating that the local prediction errors constitute a sufficient learning signal to match the generative power of TBPTT without the memory and computational overhead. Moreover, for this particular case, Predictive E-prop converged significantly faster than TBPTT, achieving the correct dynamics in 70% less epochs (*∼*23 epochs vs *∼*80 epochs). These findings hint at a possible advantage provided by the truly online nature of the training algorithm, although further studies are required to confirm the generality of these results.

**Figure 5.**
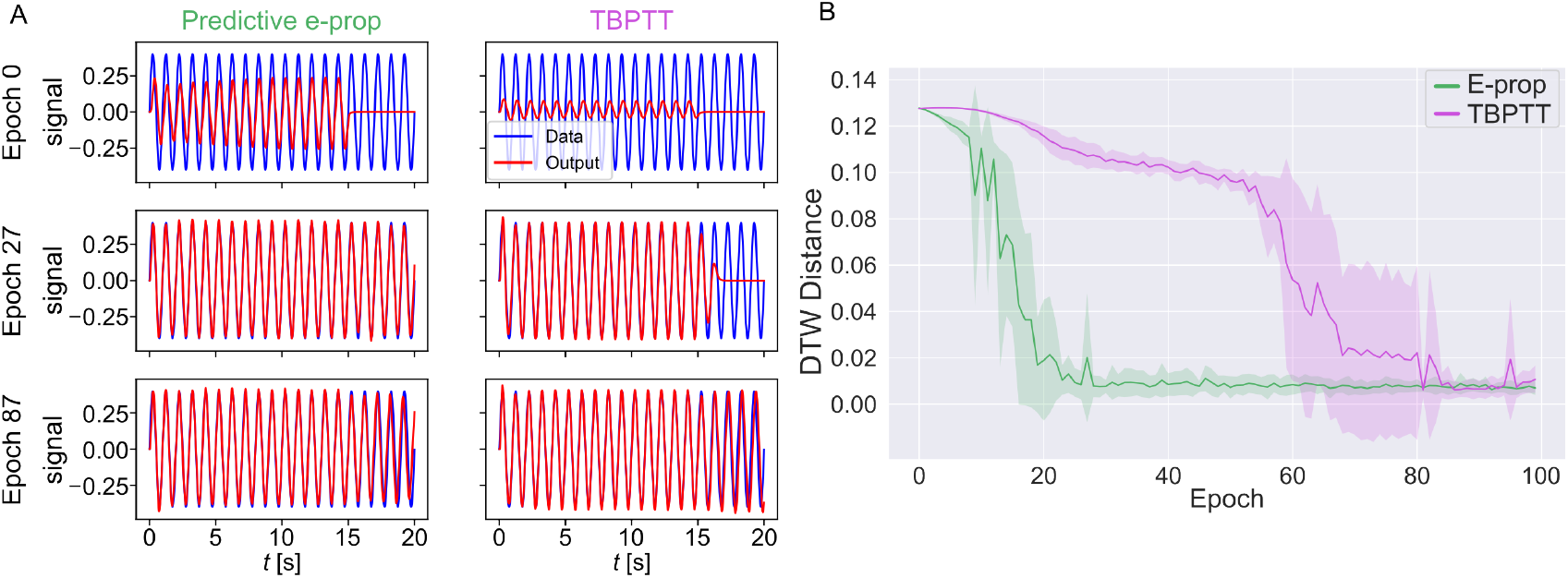
A) Examples of sine generation task using Predictive E-prop (left) and truncated BPTT (right). B) Evolution of the DTW distance for Predictive E-prop (green) and TBPTT (purple). Averages over ten randomly initialized runs are shown, with shaded areas representing the standard deviations.

## 4. Discussion

In this study, we investigated a biologically plausible approach to time-series generation, introducing Predictive E-prop: a model that combines the effectiveness of the e-prop learning algorithm [17] with the biological plausibility and efficiency inherent to the predictive coding framework [2]. Our results demonstrate that the objective of predictive coding can be optimized in RSNNs using local and online e-prop updates. Furthermore, the model proved effective in signal reconstruction tasks, showing robust denoising abilities and reconstructing sub-sampled time-series data. Finally, comparison against a TBPTT-trained analogue on the sine wave generation task revealed comparable generative performance, indicating that e-prop achieves competitive accuracy while relying solely on local information for weight updates.

Biology and neuroscience have inspired the field of artificial intelligence (AI) since its inception [40]. However, throughout its evolution, AI has strayed from its biological origins, increasing their capabilities [41, 42] but leading to two critical challenges: the unsustainable growth of human-labeled data required [43] and the ever increasing energy demand for training and inference [44]. Among many biologically inspired approaches to solving these challenges, such as the Forward-Forward algorithm [45] or heterogeneous networks [46], models based on the predictive coding framework show significant promise.

Prior works explored predictive coding using different frameworks, such as deep learning [47], rate-coding reservoirs [8, 48], and spiking reservoirs [9]. Another branch of research implemented predictive coding in RSNNs focusing on biological plausibility, employing a Hebbian-like learning rule with explicit error-detecting neurons to modulate network activity [10, 49]. The present work expands on previous predictive coding research by integrating it with e-prop [17, 20], demonstrating the feasibility of optimizing a trainable recurrent spiking layer using a gradient-based local learning rule that is compatible with the constraints of neuromorphic hardware, giving our findings particular relevance for neuromorphic engineering [23, 50, 51].

To further evaluate the suitability of Predictive E-prop for edge applications, future work should assess the performance on more realistic tasks, such as event-driven vision processing [52, 53] or power consumption prediction [54]. Moreover, in-vitro implementations of RSNNs [55, 56, 57] could provide an experimental platform to further validate the biological plausibility of the proposed model. Lastly, predictive coding was originally formulated not only as a model for perceptual inference, but also to influence the environment through motor actions [58, 59] and active inference [60] so that it adapts to the model. For instance, implementation of Predictive E-prop into reinforcement learning settings [26] could enable a closed-loop system in which actions are generated to minimize surprise. Such an extension could facilitate, e.g., adaptive sensory feedback in neuroprosthetic devices [61, 62].

## Acknowledgments

The work was partly supported by MEXT Grant-in-Aid for Transformative Research Areas (A) “Multicellular Neurobiocomputing” (24H02330, 24H02332), JSPS KAKENHI (22K19821, 22KK0177, 23H02805, 23K24913, 23K28179, 25H00447), JST ALCA-Next (JPMJAN23F3), the WISE Program for AI Electronics by Tohoku University, and the Cooperative Research Project Program of the Research Institute of Electrical Communication (RIEC) at Tohoku University. This research was partly carried out at the Laboratory for Nanoelectronics and Spintronics, RIEC, Tohoku University.

## Data availability

The data that support the findings of this study are available upon reasonable request to the authors.

## Supplementary Materials

### Supplementary Text

#### Network Regularization

In this work, a regularization term was added to the loss function to emulate the effects of homeostatic plasticity observed in biological neural networks [1]. The regularization term *E*_reg_ is defined as:

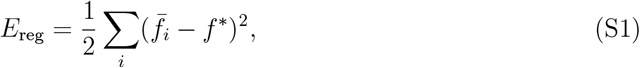

where 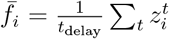 is the average firing rate of the *i*th neuron over a time span of *t*_delay_, and *f*^*∗*^ is the target firing rate which was set to 10 Hz. After *t*_delay_ ms, the average firing rate is computed and the learning signal is provided to the network.

A weight regularization term was also added to the network to mitigate excessive growth of the weights, preventing the network from falling into over-training. This term *E*_W_ is defined as:

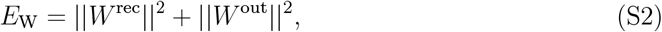

where *W* ^rec*/*out^ are the recurrent and output weights.

These additional loss terms contribute to the evolution of the network in parallel to the e-prop rule described in the main text and are incorporated into the total loss function *E* as:

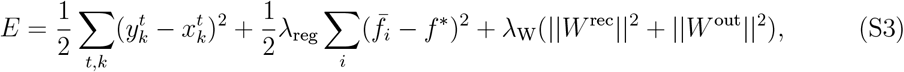

where 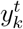 and 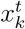 are the output of the network and the target signal at time *t* for dimension *k*, respectively. The terms *λ*_reg_ and *λ*_W_ are tunable constants that control the strength of the firing-rate and weight regularization terms, respectively.

#### Truncated BPTT

Truncated BPTT (TBPTT) [2] is an approximation of traditional backpropagation through time [3], which aims to reduce resource requirements while also allowing for online learning. The basic idea is to process sensory information for a fixed amount (*k*) of time-steps, perform backpropagation and then resume the forward process (Fig. S1A). This approximation allows the resource cost of TBPTT to become *O*(*N × k*), which is more tractable than the *O*(*N × K*) of traditional BPTT, where *K* is the total duration of the time series. The weight update rule for TBPTT is summarized as follows:

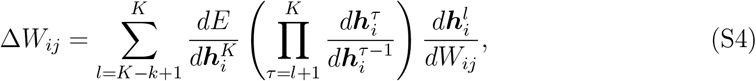

where *E* is the loss function, ***h***_*i*_ represents the hidden variables of the *i*-th neuron, and *W* is the weight matrix evolved through TBPTT. To find the optimal truncation length *k*, a parameter search was carried out, finding it to be at *k* = 100 (Fig. S1B).

## Supplementary Tables

**Table SI.** Hyperparameters for the sine wave generation experiment.

| Parameter | Value | Parameter | Value |
| --- | --- | --- | --- |
| training phase | 10000 ms | error-driven phase | 5000 ms |
| free-running phase | 5000 ms | $\eta$ | 0.0004 |
| $\lambda_W$ | 5e-06 | $\lambda_{\text{reg}}$ | 2 |
| $\gamma_d$ | 0.3 | time step ($dt$) | 1 ms |
| $t_{\text{delay}}$ | 10 ms | $f^*$ | 0.01 kHz |
| $A$ | 0.4 | $T$ | 1000 ms |
| $\phi$ | 0 | $c$ | 0.0 |
| $N_{\text{LIF}}$ | 100 | $N_{\text{ALIF}}$ | 200 |
| sparsity | 0.99 | $\tau_s$ | 250 ms |
| $\sigma_s$ | 1 | $\mu_s$ | 0 |
| $W^{\text{in}}$ | $\mathcal{N}(0, 1)/\sqrt{g}$ | $W^{\text{rec}}$ | $\mathcal{N}(0, 1)/\sqrt{N_{\text{LIF}} + N_{\text{ALIF}}}$ |
| $W^{\text{fb}}$ | $\mathcal{N}(0, 1)/\sqrt{g}$ | $W^{\text{out}}$ | $\mathcal{N}(0, 1)/\sqrt{N_{\text{LIF}} + N_{\text{ALIF}}}$ |
| $g$ | 150 | | |
| $v_{\text{th}}$ | 0.6 mV | $I_{\text{bias}}$ | 0.02 mV |
| $\tau_{\text{mem}}$ | 20 ms | $\tau_{\text{alif}}$ | 2000 ms |
| $\tau_r$ | 50 ms | $\tau_d$ | 125 ms |

**Table SII.** Hyperparameters for the Lorenz attractor generation experiment.

| Parameter | Value | Parameter | Value |
| --- | --- | --- | --- |
| training phase | 10000 ms | error-driven phase | 5000 ms |
| free-running phase | 5000 ms | $\eta$ | 0.0002 |
| $\lambda_W$ | 5e-06 | $\lambda_{\text{reg}}$ | 10 |
| $\gamma_d$ | 0.3 | time step ($dt$) | 1 ms |
| $t_{\text{delay}}$ | 10 ms | $f^*$ | 0.01 kHz |
| $x_0$ | 5 | $y_0$ | 5 |
| $z_0$ | 5 | | |
| $N_{\text{LIF}}$ | 100 | $N_{\text{ALIF}}$ | 200 |
| sparsity | 0.99 | $\tau_s$ | 250 ms |
| $\sigma_s$ | 1 | $\mu_s$ | 0 |
| $W^{\text{in}}$ | $\mathcal{N}(0, 1)/\sqrt{g}$ | $W^{\text{rec}}$ | $\mathcal{N}(0, 1)/\sqrt{N_{\text{LIF}} + N_{\text{ALIF}}}$ |
| $W^{\text{fb}}$ | $\mathcal{N}(0, 1)/\sqrt{g}$ | $W^{\text{out}}$ | $\mathcal{N}(0, 1)/\sqrt{N_{\text{LIF}} + N_{\text{ALIF}}}$ |
| $g$ | 35 | | |
| $v_{\text{th}}$ | 0.6 mV | $I_{\text{bias}}$ | 0.02 mV |
| $\tau_{\text{mem}}$ | 20 ms | $\tau_{\text{alif}}$ | 2000 ms |
| $\tau_r$ | 50 ms | $\tau_d$ | 125 ms |

**Table SIII.**
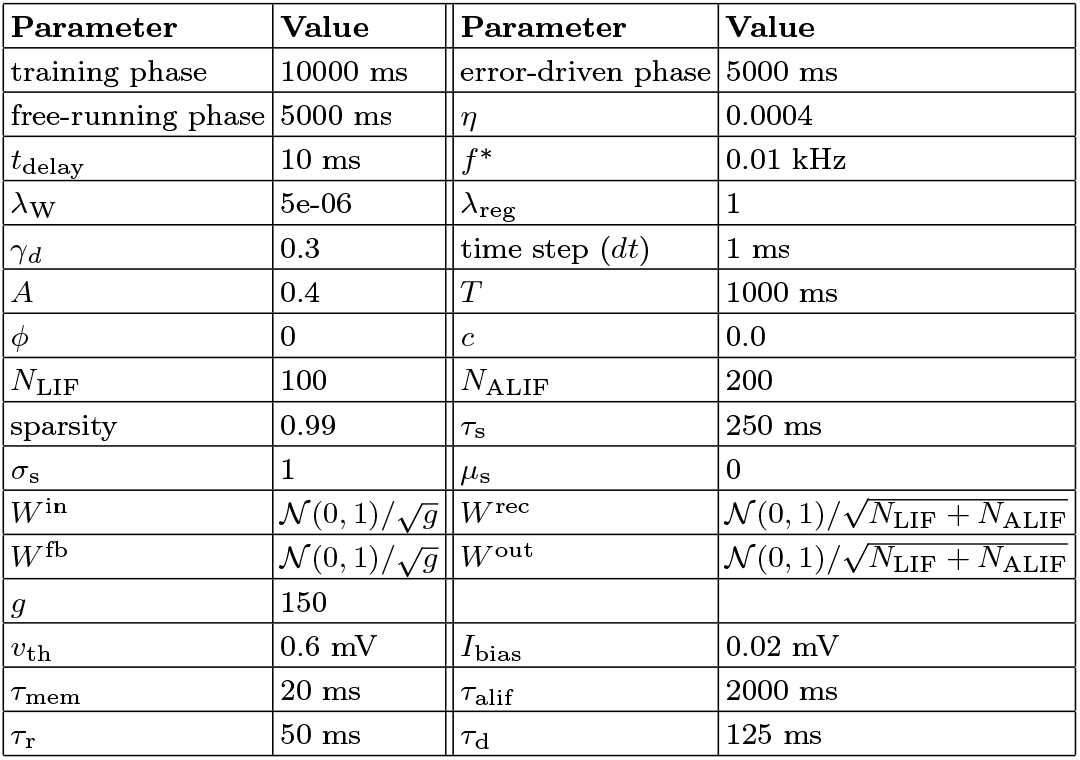
Hyperparameters for the sine wave reconstruction experiment.

**Table SIV.** Hyperparameters for the Lorenz attractor reconstruction experiment.

| Parameter | Value | Parameter | Value |
| --- | --- | --- | --- |
| training phase | 10000 ms | error-driven phase | 5000 ms |
| free-running phase | 5000 ms | $\eta$ | 0.0002 |
| $\lambda_W$ | 5e-06 | $\lambda_{\text{reg}}$ | 5 |
| $t_{\text{delay}}$ | 100 ms | $f^*$ | 0.01 kHz |
| $\gamma_d$ | 0.3 | time step ($dt$) | 1 ms |
| $x_0$ | 5 | $y_0$ | 5 |
| $z_0$ | 5 | | |
| $N_{\text{LIF}}$ | 100 | $N_{\text{ALIF}}$ | 200 |
| sparsity | 0.99 |  |  |
| $\sigma_s$ | 1 | $\mu_s$ | 0 |
| $W^{\text{in}}$ | $\mathcal{N}(0, 1)/\sqrt{g}$ | $W^{\text{rec}}$ | $\mathcal{N}(0, 1)/\sqrt{N_{\text{LIF}} + N_{\text{ALIF}}}$ |
| $W^{\text{fb}}$ | $\mathcal{N}(0, 1)/\sqrt{g}$ | $W^{\text{out}}$ | $\mathcal{N}(0, 1)/\sqrt{N_{\text{LIF}} + t_{\text{ALIF}}}$ |
| $g$ | 35 | | |
| $v_{\text{th}}$ | 0.6 mV | $I_{\text{bias}}$ | 0.02 mV |
| $\tau_{\text{mem}}$ | 20 ms | $\tau_{\text{alif}}$ | 2000 ms |
| $\tau_{\text{r}}$ | 50 ms | $\tau_{\text{d}}$ | 125 ms |

## Supplementary Figures

**Figure S1.**
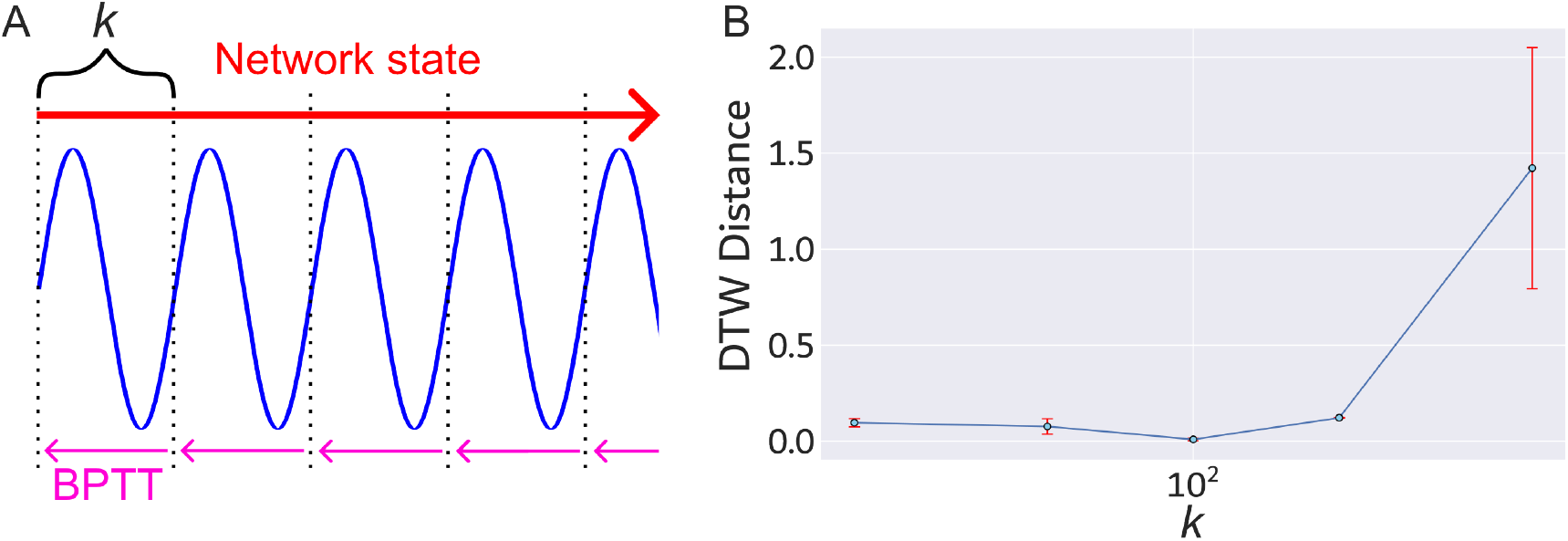
A) Schematic representation of the computational flow for TBPTT. B) Truncation length study. The optimal value found was *k* = 100.

## References

[1] G. B. Keller, T. D. Mrsic-Flogel, Predictive processing: A canonical cortical computation, Neuron 100 (2) (2018) 424–435.

[2] R. P. Rao, D. H. Ballard, Predictive coding in the visual cortex: A functional interpretation of some extra-classical receptive-field effects, Nature Neuroscience 2 (1) (1999) 79–87.

[3] A. W. N’dri, W. Gebhardt, C. Teulière, F. Zeldenrust, R. P. Rao, J. Triesch, A. Ororbia, Predictive coding with spiking neural networks: a survey, Neural Networks (2025) 108371.

[4] M. Spratling, Predictive coding as a model of biased competition in visual attention, Vision Research 48 (12) (2008) 1391–1408.

[5] M. Zhang, R. Chitic, S. M. Bohté, Energy optimization induces predictive-coding properties in a multi-compartment spiking neural network model, PLOS Computational Biology 21 (6) (2025) 1–23.

[6] A. Ali, N. Ahmad, E. de Groot, M. A. J. van Gerven, T. C. Kietzmann, Predictive coding is a consequence of energy efficiency in recurrent neural networks, Patterns 3 (12) (2022) 100639.

[7] J. Lee, J. Jo, B. Lee, J.-H. Lee, S. Yoon, Brain-inspired predictive coding improves the performance of machine challenging tasks, Frontiers in Computational Neuroscience 16 (2022) 1062678.

[8] Y. Yonemura, Y. Katori, Dynamical predictive coding with reservoir computing performs noise-robust multi-sensory speech recognition, Frontiers in Computational Neuroscience 18 (2024) 1464603.

[9] Y. Ishikawa, T. Shinkawa, T. Sumi, H. Kato, H. Yamamoto, Y. Katori, Integrating predictive coding with reservoir computing in spiking neural network model of cultured neurons, Nonlinear Theory and Its Applications, IEICE 15 (2) (2024) 432–442.

[10] A. Ororbia, Spiking neural predictive coding for continually learning from data streams, Neurocomputing 544 (2023) 126292.

[11] A. W. N’dri, T. Barbier, C. Teulière, J. Triesch, Predictive coding light, Nature Communications 16 (1) (2025) 8880.

[12] D. E. Rumelhart, G. E. Hinton, R. J. Williams, Learning representations by back-propagating errors, Nature 323 (6088) (1986) 533–536.

[13] Y. LeCun, Y. Bengio, G. Hinton, Deep learning, Nature 521 (7553) (2015) 436–444.

[14] P. J. Werbos, Backpropagation through time: What it does and how to do it, Proceedings of the IEEE 78 (10) (1990) 1550–1560.

[15] R. J. Williams, D. Zipser, A learning algorithm for continually running fully recurrent neural networks, Neural Computation 1 (2) (1989) 270–280.

[16] Y. Bengio, D.-H. Lee, J. Bornschein, T. Mesnard, Z. Lin, Towards biologically plausible deep learning, arXiv preprint arXiv:1502.04156 (2015).

[17] G. Bellec, F. Scherr, A. Subramoney, E. Hajek, D. Salaj, R. Legenstein, W. Maass, A solution to the learning dilemma for recurrent networks of spiking neurons, Nature Communications 11 (1) (2020) 3625.

[18] E. O. Neftci, H. Mostafa, F. Zenke, Surrogate gradient learning in spiking neural networks: Bringing the power of gradient-based optimization to spiking neural networks, IEEE Signal Processing Magazine 36 (6) (2019) 51–63.

[19] W. Gerstner, M. Lehmann, V. Liakoni, D. Corneil, J. Brea, Eligibility traces and plasticity on behavioral time scales: experimental support of neohebbian three-factor learning rules, Frontiers in Neural Circuits 12 (2018) 53.

[20] D. Noè, H. Yamamoto, Y. Katori, S. Sato, Efficient connectivity and intrinsic noise separation in recurrent spiking neural networks trained with e-prop, Neuromorphic Computing and Engineering 5 (4) (2025) 044002.

[21] A. Rostami, B. Vogginger, Y. Yan, C. G. Mayr, E-prop on SpiNNaker 2: Exploring online learning in spiking rnns on neuromorphic hardware, Frontiers in Neuroscience 16 (2022) 1018006.

[22] A. Perrett, S. Summerton, A. Gait, O. Rhodes, Online learning in SNNs with e-prop and neuromorphic hardware, in: Proceedings of the 2022 Annual Neuro-Inspired Computational Elements Conference, Association for Computing Machinery, 2022, p. 32–39.

[23] C. Frenkel, G. Indiveri, Reckon: A 28nm sub-mm2 task-agnostic spiking recurrent neural network processor enabling on-chip learning over second-long timescales, IEEE International Solid-State Circuits Conference (ISSCC) 65 (2022) 468–469.

[24] G. Bellec, D. Salaj, A. Subramoney, R. Legenstein, W. Maass, Long short-term memory and learning-to-learn in networks of spiking neurons, in: S. Bengio, H. Wallach, H. Larochelle, K. Grauman, N. Cesa-Bianchi, R. Garnett (Eds.), Advances in Neural Information Processing Systems, Vol. 31, 2018.

[25] W. Gerstner, W. M. Kistler, R. Naud, L. Paninski, Neuronal dynamics: From single neurons to networks and models of cognition, Cambridge University Press, 2014.

[26] D. Ha, J. Schmidhuber, Recurrent world models facilitate policy evolution, in: S. Bengio, H. Wallach, H. Larochelle, K. Grauman, N. Cesa-Bianchi, R. Garnett (Eds.), Advances in Neural Information Processing Systems, Vol. 31, 2018.

[27] G. E. Uhlenbeck, L. S. Ornstein, On the theory of the brownian motion, Physical Reviews 36 (1930) 823–841.

[28] T. P. Lillicrap, D. Cownden, D. B. Tweed, C. J. Akerman, Random synaptic feedback weights support error backpropagation for deep learning, Nature Communications 7 (1) (2016) 13276.

[29] T. Giorgino, Computing and visualizing dynamic time warping alignments in r: the dtw package, Journal of Statistical Software 31 (2009) 1–24.

[30] D. Sussillo, L. F. Abbott, Generating coherent patterns of activity from chaotic neural networks, Neuron 63 (4) (2009) 544–557.

[31] M. Lukoševičius, H. Jaeger, Reservoir computing approaches to recurrent neural network training, Computer Science Review 3 (3) (2009) 127–149.

[32] P. Vincent, H. Larochelle, Y. Bengio, P.-A. Manzagol, Extracting and composing robust features with denoising autoencoders, in: Proceedings of the 25th International Conference on Machine Learning, Association for Computing Machinery, 2008, p. 1096–1103.

[33] T. S. Lee, M. Nguyen, Dynamics of subjective contour formation in the early visual cortex, Proceedings of the National Academy of Sciences 98 (4) (2001) 1907–1911.

[34] R. Von der Heydt, E. Peterhans, G. Baumgartner, Illusory contours and cortical neuron responses, Science 224 (4654) (1984) 1260–1262.

[35] R. Näätänen, A. W. Gaillard, S. Mäntysalo, Early selective-attention effect on evoked potential reinterpreted, Acta Psychologica 42 (4) (1978) 313–329.

[36] H. Brown, R. A. Adams, I. Parees, M. Edwards, K. Friston, Active inference, sensory attenuation and illusions, Cognitive Processing 14 (4) (2013) 411–427.

[37] I. Sutskever, Training recurrent neural networks, Ph.D. thesis, University of Toronto (2013).

[38] R. Pascanu, T. Mikolov, Y. Bengio, On the difficulty of training recurrent neural networks, in: International Conference on Machine Learning, PMLR, 2013, pp. 1310–1318.

[39] R. J. Williams, J. Peng, An efficient gradient-based algorithm for online training of recurrent network trajectories, Neural Computation 2 (4) (1990) 490–501.

[40] F. Rosenblatt, The perceptron: A probabilistic model for information storage and organization in the brain., Psychological Review 65 (6) (1958) 386.

[41] A. Vaswani, N. Shazeer, N. Parmar, J. Uszkoreit, L. Jones, A. N. Gomez, L. Kaiser, I. Polosukhin, Attention is all you need, Advances in Neural Information Processing Systems 30 (2017) 5998–6008.

[42] I. J. Goodfellow, J. Pouget-Abadie, M. Mirza, B. Xu, D. Warde-Farley, S. Ozair, A. Courville, Y. Bengio, Generative adversarial nets, in: Z. Ghahramani, M. Welling, C. Cortes, N. Lawrence, K. Weinberger (Eds.), Advances in Neural Information Processing Systems, Vol. 27, 2014.

[43] Y. Roh, G. Heo, S. E. Whang, A survey on data collection for machine learning: A big data - AI integration perspective, IEEE Transactions on Knowledge and Data Engineering 33 (4) (2021) 1328–1347.

[44] S. Luccioni, Y. Jernite, E. Strubell, Power hungry processing: Watts driving the cost of ai deployment?, Proceedings of the ACM Conference on Fairness, Accountability, and Transparency (FAccT) (2024) 85–99.

[45] G. Hinton, The forward-forward algorithm: Some preliminary investigations, arXiv preprint arXiv:2212.13345 2 (3) (2022) 5.

[46] B. Wang, Y. Zhang, H. Li, H. Dou, Y. Guo, Y. Deng, Biologically inspired heterogeneous learning for accurate, efficient and low-latency neural network, National Science Review 12 (1) (2025) nwae301.

[47] B. Millidge, A. Tschantz, C. L. Buckley, Predictive coding approximates backprop along arbitrary computation graphs, Neural Computation 34 (6) (2022) 1329–1368.

[48] Y. Yonemura, Y. Katori, Network model of predictive coding based on reservoir computing for multi-modal processing of visual and auditory signals, Nonlinear Theory and Its Applications, IEICE 12 (2) (2021) 143–156.

[49] M. Lan, X. Xiong, Z. Jiang, Y. Lou, Pc-snn: Supervised learning with local hebbian synaptic plasticity based on predictive coding in spiking neural networks, arXiv preprint arXiv:2211.15386 (2022).

[50] S. Moriya, M. Ishikawa, S. Ono, H. Yamamoto, Y. Yuminaka, Y. Horio, J. Madrenas, S. Sato, Analog VLSI implementation of subthreshold spiking neural networks and its application to reservoir computing, IEEE Transactions Circuits and Systems I 72 (10) (2025) 5571–5582.

[51] B. Rajendran, A. Sebastian, M. Schmuker, N. Srinivasa, E. Eleftheriou, Low-power neuromorphic hardware for signal processing applications: A review of architectural and system-level design approaches, IEEE Signal Processing Magazine 36 (6) (2019) 97–110.

[52] G. Gallego, T. Delbrück, G. Orchard, C. Bartolozzi, B. Taba, A. Censi, S. Leutenegger, A. J. Davison, J. Conradt, K. Daniilidis, et al., Event-based vision: A survey, IEEE Transactions on Pattern Analysis and Machine Intelligence 44 (1) (2020) 154–180.

[53] P. Lichtsteiner, T. Posch, Christoph andG Delbruck, A 128 × 128 120 db 15µ s latency asynchronous temporal contrast vision sensor, IEEE Journal of Solid-State Circuits 43 (2) (2008) 566–576.

[54] S. Vazquez, J. Rodriguez, M. Rivera, L. G. Franquelo, M. Norambuena, Model predictive control for power converters and drives: Advances and trends, IEEE Transactions on Industrial Electronics 64 (2) (2016) 935–947.

[55] Y. Yada, S. Yasuda, H. Takahashi, Physical reservoir computing with force learning in a living neuronal culture, Applied Physics Letters 119 (17) (2021) 173701.

[56] T. Sumi, H. Yamamoto, Y. Katori, K. Ito, S. Moriya, T. Konno, S. Sato, A. Hirano-Iwata, Biological neurons act as generalization filters in reservoir computing, Proceedings of the National Academy of Sciences of the U.S.A 120 (25) (2023) e2217008120.

[57] Y. Sato, H. Yamamoto, Y. Ishikawa, T. Sumi, Y. Sono, S. Sato, Y. Katori, A. Hirano-Iwata, In silico modeling of reservoir-based predictive coding in biological neuronal networks on microelectrode arrays, Japanese Journal of Applied Physics 63 (10) (2024) 108001.

[58] K. Friston, The free-energy principle: A unified brain theory?, Nature Reviews Neuroscience 11 (2) (2010) 127–138.

[59] H. Brown, K. Friston, S. Bestmann, Active inference, attention, and motor preparation, Frontiers in Psychology 2 (2011) 218.

[60] K. J. Friston, J. Daunizeau, S. J. Kiebel, Reinforcement learning or active inference?, PLOS One 4 (7) (2009) e6421.

[61] C. Pezzato, R. Ferrari, C. H. Corbato, A novel adaptive controller for robot manipulators based on active inference, IEEE Robotics and Automation Letters 5 (2) (2020) 2973–2980.

[62] A. Zangrandi, M. D’Alonzo, C. Cipriani, G. Di Pino, Neurophysiology of slip sensation and grip reaction: Insights for hand prosthesis control of slippage, Journal of Neurophysiology 126 (2) (2021) 477–492.

## References

[1] G. G. Turrigiano, S. B. Nelson, Homeostatic plasticity in the developing nervous system, Nat. Rev. Neurosci. 5 (2) (2004) 97–107.

[2] I. Sutskever, Training recurrent neural networks, Ph.D. thesis, University of Toronto (2013).

[3] P. J. Werbos, Backpropagation through time: What it does and how to do it, Proceedings of the IEEE 78 (10) (1990) 1550–1560.

